# Recording animal-view videos of the natural world

**DOI:** 10.1101/2022.11.22.517269

**Authors:** Vera Vasas, Mark C. Lowell, Juliana Villa, Quentin D. Jamison, Anna G. Siegle, Pavan Kumar Reddy Katta, Pushyami Bhagavathula, Peter G. Kevan, Drew Fulton, Neil Losin, David Kepplinger, Shakiba Salehian, Rebecca E. Forkner, Daniel Hanley

## Abstract

Plants, animals, and fungi display a rich tapestry of colors. Animals, in particular, use colors in dynamic displays performed in spatially complex environments. In such natural settings, light is reflected or refracted from objects with complex shapes that cast shadows and generate highlights. In addition, the illuminating light changes continuously as viewers and targets move through heterogeneous, continually fluctuating, light conditions. Although traditional spectrophotometric approaches for studying colors are objective and repeatable, they fail to document this complexity. Worse, they miss the temporal variation of color signals entirely. Here, we introduce hardware and software that provide ecologists and filmmakers the ability to accurately record animal-perceived colors in motion. Specifically, our Python codes transform photos or videos into perceivable units (quantum catches) for any animal of known photoreceptor sensitivity. We provide the plans, codes, and validation tests necessary for end-users to capture animal-view videos. This approach will allow ecologists to investigate how animals use colors in dynamic behavioral displays, the ways natural illumination alters perceived colors, and other questions that remained unaddressed until now due to a lack of suitable tools. Finally, our pipeline provides scientists and filmmakers with a new, empirically grounded approach for depicting the perceptual worlds of non-human animals.

## Introduction

How do animals see the world? This simple question has spurred discovery since the advent of modern science (Kelber and Osorio 2010) and inspired breathtaking media illustrating what animals see. Color vision is vital for wide swaths of life (Cuthill et al. 2017). As a result, animals have adapted photoreceptors that are sensitive to a wide array of light that is not necessarily perceivable to humans. For example, many animals can detect ultraviolet or polarized light invisible to us (Chiou et al. 2008), while others can even detect infrared light, using novel sense organs (Chiou et al. 2008; Gracheva et al. 2010; Panzano et al. 2010). The consequence is that each animal sees color differently (Bennet and Thery 2007), which makes false color imagery of animal vision powerful and compelling (Eisner et al. 1969; Kevan 1972; Verhoeven et al. 2018). Unfortunately, current techniques are unable to quantify perceived colors of organisms in motion, even though such movement is often crucial for color appearance and signal detection (Rosenthal 2007; Hutton et al. 2015). Overcoming this serious barrier should spur widespread advancements in the field of sensory ecology (Cuthill et al. 2017; Tan and Elgar 2021). Here we provide a solution to these challenges in the form of a tool for researchers or filmmakers to record videos that accurately represent animal-perceived colors.

Accurately portraying animal-perceived colors requires careful consideration of the perceptual abilities of the relevant receivers (Goldsmith 1990; Chittka et al. 1994; Kelber et al. 2003). Traditional spectrophotometric approaches estimate visual stimulation by measuring an animal’s photoreceptor response to object-reflected light (Endler 1993a; Osorio and Vorobyev 2008). The reflected light, in turn, can be estimated as a function of the illuminating light and the spectral reflectance properties of the object (Vorobyev and Osorio 1998). Though accurate, individually measuring reflectance spectra for each object is laborious and all spatial and temporal information is lost in the process. Moreover, measurements must be taken from sufficiently large, uniformly colored, and relatively flat and smooth surfaces (Montgomerie 2006). Yet, many natural colors pose technical challenges: powders and latticed materials, or objects that are iridescent, glossy, transparent, translucent, or luminescent, or animals that shift their colors using iridophores (Meadows et al. 2011; Stavenga et al. 2011; Siddique et al. 2015; Teyssier et al. 2015; Ben Dor et al. 2015; Dunning et al. 2019).

Multispectral photography offers a different approach (Stevens et al. 2007; Troscianko and Stevens 2015; Vasas et al. 2017; van den Berg et al. 2019). It relies on taking a series of photos in wavelength ranges that are generally broader than standard ‘human-visible’ photographs. Typically, subjects are photographed using a camera sensitive to broadband light, through a succession of narrow-bandpass filters (Kevan 1972, 1978; Kevan and Backhaus 1998). In this way, researchers acquire images in regions surpassing human visual experience (e.g., the ultraviolet and infrared ranges), organize these as a stack of multiple, clearly differentiated color channels (Kevan 1978; Westland and Ripamonti 2004; Stevens et al. 2007), from which they can derive camera-independent measurements of color (Troscianko and Stevens 2015; Vasas et al. 2017; Tedore and Nilsson 2019). These approaches have a rich tradition in the study of pollinator-plant relations, where the relationship between colorimetry and the visual systems of diverse organisms has long been embraced (Kevan 1972, 1978, Kevan & Backhaus 1998, Chittka 1992, Verhoeven et al. 2018). The resultant multispectral images trade-off some accuracy for tremendous improvements to securing spatial information, which is often a welcomed compromise when the goal is to understand animal signals. However, by its nature the method works only on still objects, and it is unsuitable for studying the temporal aspects of signals (Rosenthal 2007).

Yet temporal changes can be central to a biological signal (Rosenthal 2007; Hutton et al. 2015). A recently proposed approach for studying dynamic color signals combines multispectral imaging with digital 3D modeling (Miller et al. 2022). The next step will be to animate the resulting 3D multispectral models and further analyze them *in silico*, simulating a variety of receiver-specific visual models or viewing conditions (Kim et al. 2012, Medina et al. 2020, Miller et al. 2022). Such models will likely play a crucial role in our understanding of the ways animal posture and the receivers’ viewpoints alter visual signals. It is both a strength and limitation that such approaches reproduce the color without the confounding factors of viewing angle or variable illumination. Yet in realistic, natural settings, illumination is directional, and the light natural stimuli reflect often changes with the viewing angle (Hutton et al. 2015). As a result, these methods will not account fully for the ways perceived colors vary in nature.

In this paper we take a different approach. We present a camera system and an associated computational framework that produces animal-view videos, with sufficient precision to be used for scientific purposes. This new tool captures the full complexity of visual signals, as perceived under natural contexts, where moving targets may be unevenly illuminated. We address these challenges by developing a camera system that records videos in four channels simultaneously (UV, blue, green and red). Data from this, or similar, systems can then be processed using a set of transformations that directly translates the video recordings to reliable estimates of perceivable units (quantum catches) for any animal of known photoreceptor sensitivity. Thus, for the first time, it is possible to record moving stimuli – crucially, living and behaving animals – and study the temporal component of visual signals.

## Results & Discussion

Our new pipeline combines existing methods of multispectral photography with a newly developed hardware design and a set of transformation functions implemented in Python to record and process videos (Figure 1). The hardware includes a beam splitter that separates ultraviolet from visible light, and directs images from these respective wavebands to two independent cameras. An advantage of this approach is that all color channels are recorded simultaneously, which enables video recording. The recordings are then linearized and transformed into animal perceived colors, in the form of photoreceptor quantum catches. Then, these animal-view videos can be further analyzed or combined to illustrate animal-perceived colors using false color imagery (Kevan 1978; Vasas et al. 2017; Verhoeven et al. 2018).

**Figure 1.**
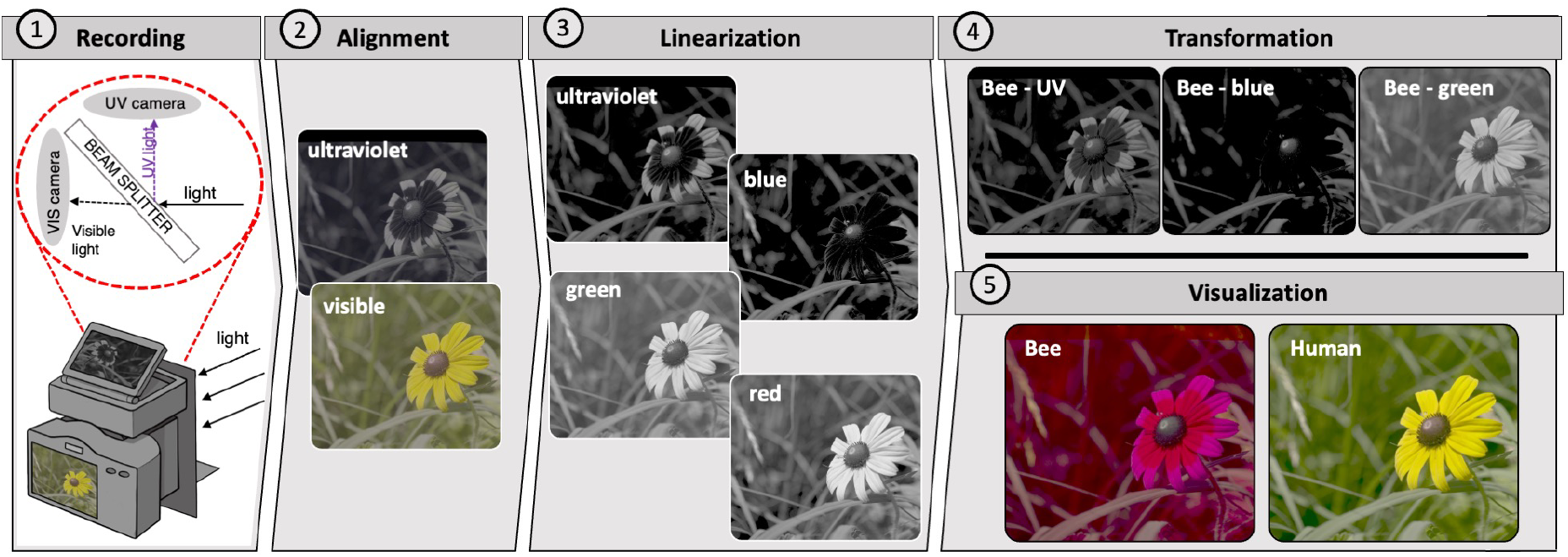
Recording and video processing pipeline. Scenes are 1) projected to an internal beam splitter that reflects ultraviolet light and passes visible light to two independent cameras. This design eliminated the need for switching filters and so allows for the rapid collection of multispectral recordings (videos or images). Then, users can use our pipeline to 2) align the recordings automatically (see Materials and Methods, Figure S8). These recordings are 3) linearized automatically using the custom color card or a set of grayscales of known reflectivity, resulting in camera quantum catches (CC). Finally, the linearized recordings are 4) transformed to animal quantum catches (AC, in this case representing honeybee *Apis mellifera* vision), which can subsequently be 5) visualized as false color images or videos (labeled as ‘bee’) and compared to the composition of the linear images or videos (labeled as ‘human’). In this case, we illustrate the pipeline using a black-eyed Susan *Rudbeckia hirta*, and display the linear and transformed images applying a gamma correction (CC_i_^0.3^ and AC_i_^0.3^, respectively). The resultant images 5) are also available in a larger format (Figure S9).

This approach accurately produces photoreceptor quantum catches for honeybees *(Apis mellifera)* or the average ultraviolet-sensitive avian viewer (Endler and Mielke 2005); however, it can work for any organisms provided users supply data on photoreceptor sensitivity. Specifically, the technique reliably captures animal perceived colors of known standards (Figure 2 and Figures S1-S6; 0.928 < R^2^ < 0.992, depending on the photoreceptor type and the illumination) and natural objects such as flowers, leaves, birds’ eggs, and feathers (Figure S7, 0.826 < R^2^ < 0.940, depending on the photoreceptor type). These surfaces constitute different color production mechanisms (Stoddard and Prum 2011, van der Kooi et al. 2016) and were photographed under either artificial illumination indoors or natural sunlight outdoors, which highlights the broader applicability of our method. Moreover, our pipeline was designed with usability in mind. As such, we have included easily accessible, consumer-level cameras, and have automated the detection of custom color standards and the linearization and transformation processes (Figure 1, Figure S8, Table S8). As part of our approach, we provide methods for quantifying estimation error. Finally, our pipeline is very flexible, allowing the user to easily swap camera systems or project to different animal systems. As all components are open source, the tools invite continued improvements (see Data availability).

**Figure 2.**
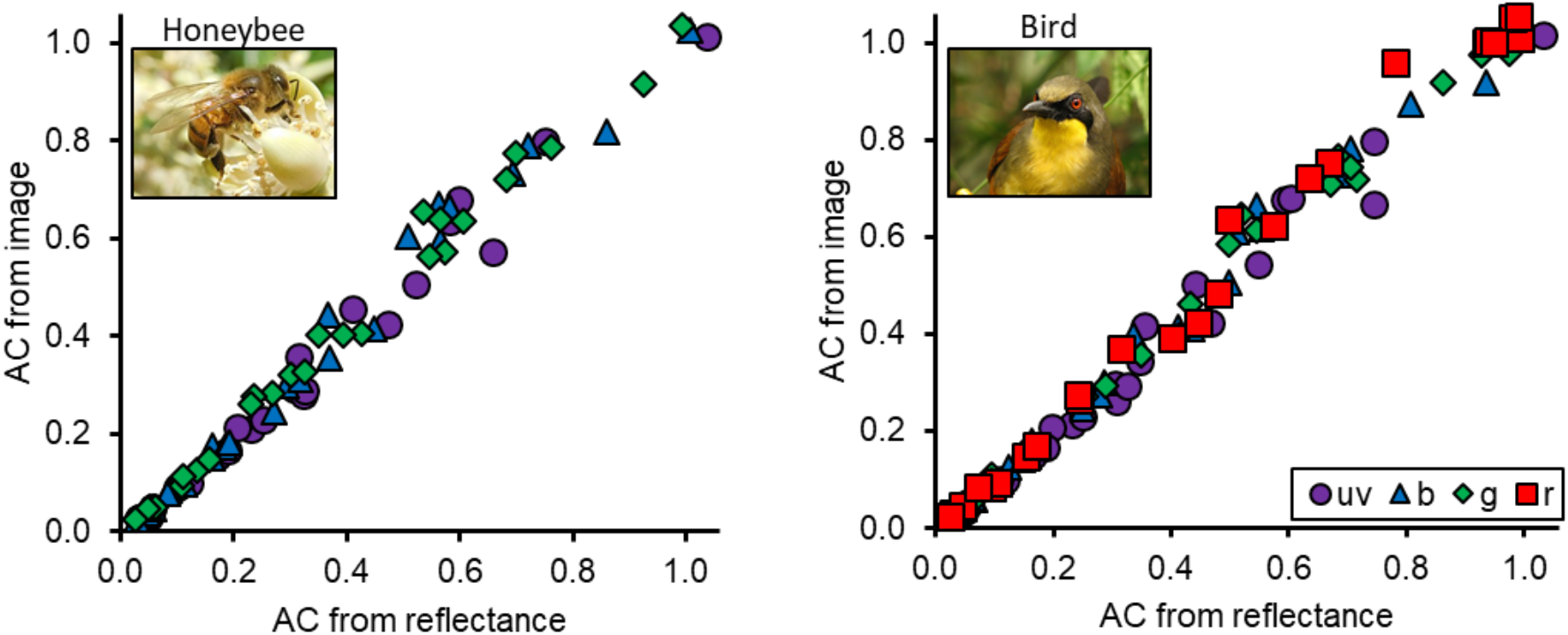
Quantum catch recovery from animal-view video. Video recordings can produce accurate estimates of animal perceived colors. In this case, we illustrate the relationship by comparing animal catches derived from our pipeline (‘AC from image’) versus animal catches derived from reflectance spectra (‘AC from reflectance’), for the honeybee (left) and average ultraviolet sensitive bird (right). We present quantum catch estimates for photoreceptors broadly sensitive to ultraviolet (violet circles), blue (blue triangle), green (green diamonds) and red light (red squares; for the bird alone). For a detailed description of the data, see Figure S5 and Table S5; for further tests see Figures S1-S4, S6-7 and Tables S1-S4, S6-7. Photo credit: honeybee *Apis mellifera* by Filo gèn’ and bird, rufous-vented laughingthrush *Pterorhinus gularis*, by Rohit Naniwadekar, both unmodified and shared under the <u>Creative Commons Attribution-Share Alike 4.0</u> license.

Animal-view videos provide a new possibility for researchers to study spatially and temporally complex, multimodal displays, in nature where these signals are produced and perceived (Figure 3). This will usher in a new age of sensory ecology. We highlight three such frontiers. First, as the method does not require moving parts (e.g. filters or lenses) common in other applications of multispectral photography, it allows for more rapid data acquisition (Figure 3d). This can speed up the pipelines of color estimation and photogrammetry projects (Miller et al. 2022; Medina et al. 2020). Second, signals and displays can be studied in their natural context. For example, in natural settings the intensity and spectral distribution of light continuously shifts and some habitats (e.g., forests) can have very patchy illumination. In addition, the incidence angle of directional light sources, such as sunlight, will alter how animals perceive objects. This effect is particularly evident on glossy or iridescent surfaces (e.g., some feathers, Figure 3d), but ultimately applies to all natural surfaces. As a result, the perceived color of a leaf fluttering in the wind or a bird walking in the undergrowth will constantly change. Our goal is to provide a simple pipeline to easily capture and analyze colorful signals as they would be perceived in the environment where they are produced and experienced by free-living organisms (*sensu* Endler 1993b; Endler and Mielke 2005). Finally, and perhaps most importantly, it is now possible to study the interplay of colors over time. By analyzing animal-view videos of the natural world, researchers can now explore a wide range of cues and signals in motion. These run the gamut from foraging and navigation cues through camouflage patterns to the intricacies of visual communication (Figure 3a-c), opening many new research avenues.

**Figure 3.**
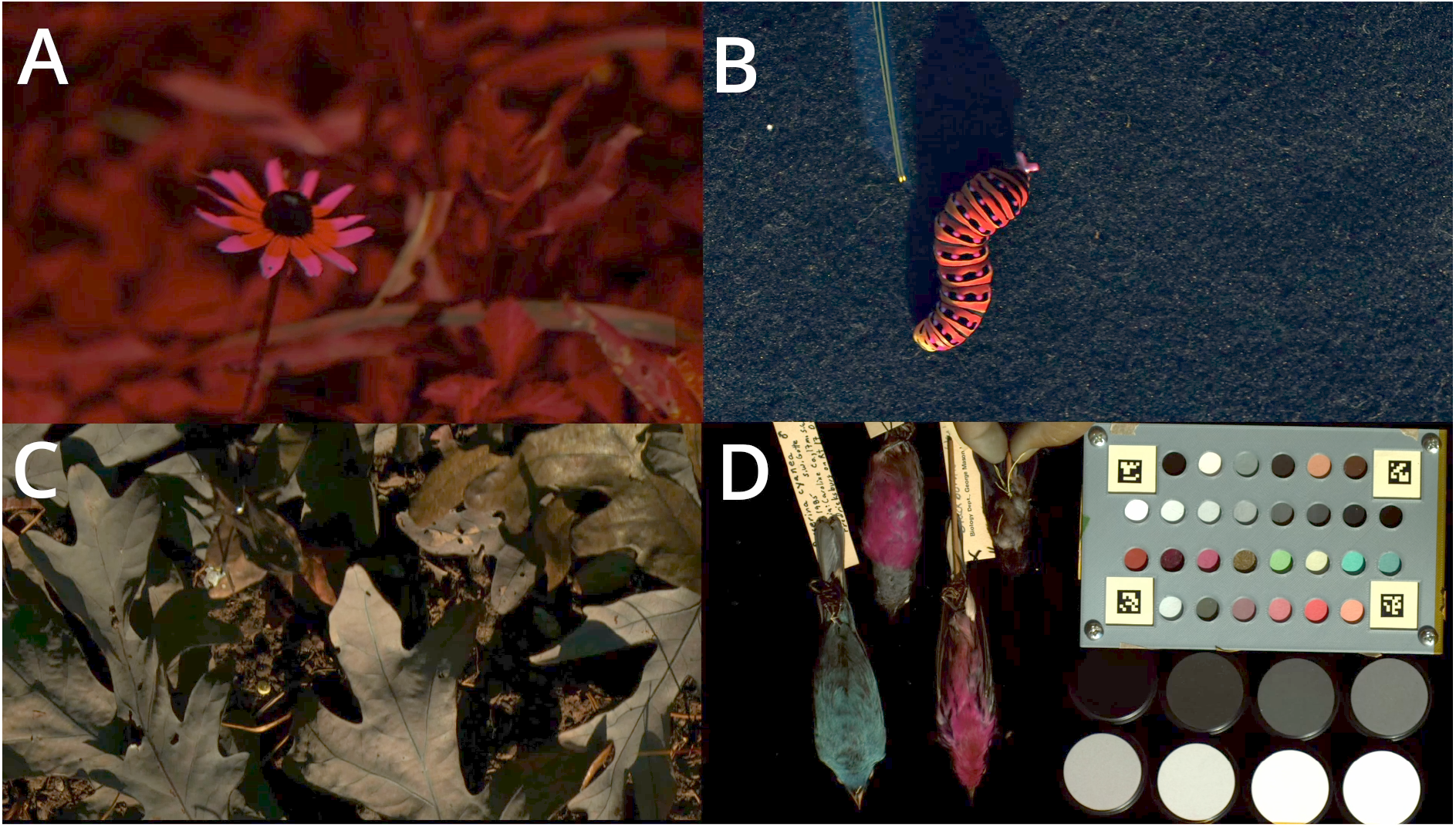
Frame excerpts from false color videos. Our pipeline can be used to produce animal-view videos of A) stimuli in motion, B) startle displays, C) camouflage in motion, and D) pigmentary and iridescent colors. Here we depict, A) the nectar guides of a black-eyed Susan *Rudbeckia hirta* under natural light as it moves in the wind. Next, we illustrate a B) black swallowtail caterpillar *Papilio polyxenes* revealing its antenna, which are otherwise concealed and would be impossible to measure using spectroscopy or traditional multispectral imaging. Likewise, our pipeline can be used to study animals such as the C) North American wheel bug *Arilus cristatus*, which is generally camouflaged and can use its motion or motion in its background to avoid detection. Finally, the pipeline provides a workflow for D) rapid data acquisition for color estimates of museum specimens, for both constant pigment-based colors or structurally-based iridescent colors. Here we illustrate frames from recordings processed as honeybee *Apis mellifera* quantum catches. To emphasize the contrast, we applied a gamma correction (AC_i_^0.5^) to this frame. The <u>video is available here</u>.

## Acknowledgements

We thank the National Geographic Society for funding (NGS-64350T), and the Office of Research, Innovation, and Economic Impact at George Mason University and Blandy Experimental Farm for additional funding. We are grateful for the support of the Smithsonian Mason School of Conservation and the George Mason’s Office of Student Scholarship, Creative Activities. We are grateful to Dat Vo, Anirudh Sai Surya Krovvidi, Marcello Tripoli, and Simran Kohli for their assistance. We also thank Dr. Alice Bridges for her help in preparing Figure 1.

## Materials and Methods

### Camera system

The camera system consists of readily available parts and cameras (Sony a6400) in a modular 3D printed housing (Figure 1a, Figure S10). The front panel supports a single camera lens (80mm f/5.6 EL-Nikkor enlarger lens) which passes light to a dichroic beam splitter (DMLP425R, Thorlabs), placed at a 45° angle relative to the beam path. This beam splitter reflects short wavelength light (<425nm), which, after passing through a shortpass filter (FF01-390/25, Semrock) that only transmits short wavelength light (<390nm), reaches a full spectrum camera. Our full spectrum camera was modified by Lifepixel (https://www.lifepixel.com/); however, a range of vendors offer this service (Crowther 2019). Simultaneously, visible light (~425 nm - 720nm) passes through the beamsplitter and is captured by a stock camera at the rear of the housing. Both cameras are triggered simultaneously using a spliced cable release. To allow dynamic focusing, the front lens mounting plate is attached to a sliding rail with a bag bellows placed between the lens and housing.

The system is designed to be flexible. The 3D printed housing allows users to design new mounting plates, typically just a single side of the cube-shaped housing, to allow for different lenses, cameras, or even internal optical components (e.g., the beam splitter or shortpass filter). For example, while we have built our system around capturing ultraviolet and human-visible portions of the light spectrum, this system could easily be adapted to capture infrared by swapping out the filter, beam splitter, and (possibly) the lens. In its simplest form, the lens can be fixed directly to the housing which results in a fixed focal plane. We used a sliding rail to allow dynamic focusing, but if space allows, a focusing helicoid can be attached to the front lens mounting plate. All parts of this model are available for download (see Data availability).

### Shooting images and videos with the camera system

It is important to note that the angles of the camera, subject, and illuminant will have notable and potentially unexpected effects on the color (Osorio and Ham 2002). Because our goal is to capture colors as perceived, the camera needs to be positioned at the vantage point of the relevant animal receiver (e.g., overhead or from ground level, depending on the target species). Users should also carefully consider where to shoot and how the camera (animal viewer) should be positioned, avoiding obvious situations where subjects are backlit (e.g., viewed against a bright sky). Both cameras should be set to record using S-Log3, which applies a specific gamma function that reduces the likelihood of under- or overexposure. As a result, the S-Log3 format provides greater access to the camera’s full dynamic range (facilitating color grading). RGB values recorded in this way closely correspond with the camera’s raw sensor responses for both jpg and mp4 files (Sony Corporation 2016). SONY provides transformation functions to transform the S-Log3 to native ‘reflectance’ (Sony Corporation 2016), but these are also recoverable using either a polynomial or power function (see *Linearization* below), as described elsewhere (Westland and Ripamonti 2004; Stevens et al. 2007).

Once in position the camera’s shutter speed can be adjusted such that a set of isoluminant (spectrally flat) grayscale (white to black) standards is properly exposed. We recommend a custom grayscale made from mixing barium sulfate paint (Labsphere 6080) with a spectrally flat black paint (Culture Hustle, Black 3.0), which is reliable and (comparatively) inexpensive. Alternatively, a commercially available set of eight calibrated Spectralon standards reflecting 99, 80, 60, 40, 20, 10, 5, and 2% of incident light (Labsphere, RSS-08-020) produces excellent results. In this study, we use both custom and Spectralon grayscales. The calibration needs to be repeated for each light condition. When recording images, the calibration shots should be taken at least once for a photo session, but taking calibration shots periodically, or including the standards in each image is ideal. When recording videos, the calibration frames should be included the video. However, just as with other approaches (Stevens et al. 2007; Troscianko and Stevens 2015), the calibration can also be completed just after photographing or filming a behavioral display as long as the lighting conditions have not changed.

To validate the accuracy of the transformation (see *Transformation* and *Validation tests* below), it is important to record colors that have substantial variation in reflectance over a broad spectral range encompassing the visual sensitivity of the target organism (e.g., 300-700 nm). As in other studies (Troscianko and Stevens 2015), we used commercially available pastels (Blick Art Supplies, 20 Artists’ Pastels Half Sticks. 21948-1209) alongside the previously described custom grayscale made from a barium sulfate and flat black paint (Figure S8). These color and grayscale standards were held within a 3d printable holder made from gray PLA plastic with ~50% reflectance across the 300-700 nm range (Prusa, PLA Silver), with four Augmented Reality University of Cordoba (ARUCO) fiducial markers (Garrido-Jurado et al., 2014) used for automatic alignment, linearization, and validation tests (see below).

### Estimating sensor sensitivity

We used a monochromator to measure the sensitivity of the camera sensors from 280 to 800 nm (Hubel et al. 1994; Normand et al. 2007; Sigernes et al. 2009; Crowther et al. 2019). Specifically, we connected a xenon light source (SLS 205, Thorlabs) to a monochromator (Optimetrics, DMC1-03) via a 1000μm single fiber (58458, Edmund Optics). Then, light from the monochromator was passed through a second 1000μm single fiber (58458, Edmund Optics) and collimating lens (Ocean Optics, 74-ACH) to a radiometrically calibrated spectrometer (Ocean Optics, Jaz) with a cosine corrector made of Spectralon (CC-3-DA cosine). We incrementally varied light from 280nm to 800nm in ~5nm increments in relatively narrow bands (mean FWHM ± s.e. = 7.3 ± 0.29 nm, Figure S11). At each increment, we measured the absolute irradiance at a consistent distance and orientation and simultaneously we photographed the illuminated cosine corrector using our camera system in RAW (ARW) format, setting the visible camera’s ISO to 100 and the ultraviolet camera’s ISO to 2000. Then, we calculated sensitivity as

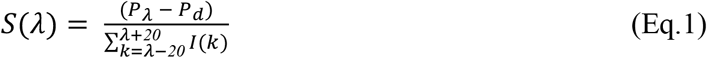

Where S is the sensor sensitivity,*P_λ_* is the pixel value for each measurement area (i.e., the area that is illuminated), *P_d_* is the sensor value for the image from an equivalent sized dark portion of the frame (i.e., dark signal), and *I* is the irradiance of the light projected on the radiometer converted to photon flux (μmol s^-1^ m^-2^). In this case, we used the sum photon flux around the main peak to reduce the impact of second order scattering, which was detected at wavelengths ≥721nm (second order peaks at ≥360.5nm). We collected a second set of measurements on the ultraviolet camera where we inserted a longpass filter (415nm, MidOpt LP415/25) within the beam path to block second order scatter (this reduced the pixel values of the UV sensor from 39.3 ± 2.7 to 1.7 ± 0.05). This confirmed that our sensitivity estimates were not influenced by second order scatter and that the ultraviolet camera was insensitive to longer wavelength light (mean ± se difference from dark pixel = 0.14 ± 0.05). Finally, each sensor was relativized by dividing the estimated sensitivity at each wavelength by the total (sum) sensitivity, such that the sum of the resultant sensor sensitivity was equal to one for each sensor (Figure S12).

### Alignment

An initial step of the pipeline involves a two-stage process of aligning the visual and ultraviolet cameras. First, we apply a manually-designed coarse alignment of the UV channel consisting of a flip, rotation, translation, and crop. This initial coarse alignment can be reused because the cameras’ relative position remains consistent. Although our modular system holds both cameras securely, very slight shifts between sessions are inevitable, so as a second step we calculate a fine-grained correction to the coarse alignment. If the ARUCO fiducial markers are visible in an image pair, a homography warp can be calculated by localizing the corners of the markers. Alternatively, we can use the Enhanced Correlation Coefficient (ECC) algorithm (Evangelidis and Psarakis, 2008). Since these algorithms can sometimes fail when image conditions are poor, we calculate several candidate alignments using different image pairs. The number of pairs used is determined by how many images can be stored in memory at a time. Then we apply each candidate alignment to a large batch of image pairs and evaluate them using the mean ECC across each image pair, selecting the alignment with the best average ECC. For videos, the frames also need to be aligned temporally, to ensure that the frames recorded by each camera correspond with one another. We calculate the spatial alignment for a range of temporal offsets, then select the best offset by calculating the mean ECC value just as we do when selecting the optimal spatial alignment.

### Linearization

Many digital cameras do not directly output raw video recordings or these are impractical for applications in the field. Rather, sensor responses are typically subjected to a variety of conversions, such as Gamma correction and white balancing, to produce images optimized for a human viewer (Westland and Ripamonti 2004). Therefore, a necessary prerequisite for converting these data to animal photoreceptor quantum catches is to estimate the relative photon capture of the camera system’s sensors (hereafter, camera quantum catches). To do so we generate linearized sensor output from each of the camera’s color channels (Westland and Ripamonti 2004; Pike 2011). This is achieved by empirically finding the relationship between recorded and ideal linearized pixel values.

The recorded pixel values should refer to a set of isoluminant standards with known reflectance curves. In this study we used a custom grayscale (Figure S8) made from mixing barium sulfate paint (Labsphere 6080) with a spectrally flat black paint (Culture Hustle, Black 3.0), and a commercially available set of eight calibrated Spectralon standards reflecting 99, 80, 60, 40, 20, 10, 5, and 2% of incident light (Labsphere, RSS-08-020). To reduce the time required for manually selecting calibration and test samples within the images, we provide an automated process that reads the four ARUCO fiducial markers (Garrido-Jurado et al., 2014) on our custom color standard and extracts the pixel values of the custom standards. The ARUCO algorithm provides a fast, highly accurate, automated method to locate relevant points on the custom color card.

We calculate the ideal linearized values by predicting the expected camera quantum catch under ideal light conditions for a set of isoluminant standards. Specifically, we calculate the camera catch that would be expected under ideal (isoluminant) illumination for each channel as.

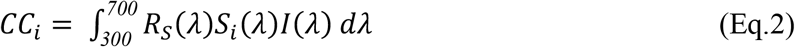

where *CC_i_* is the camera quantum catch of the sensor *i*, *R_s_* is the spectral reflectance function of the stimulus, *S_i_* is the spectral sensitivity function of the sensor, and I is the photon flux of the illuminant, in this case set to 1 across all wavelengths.

Having established the target values in this way, we then fit either a power law or a polynomial regression for each channel using the trust region reflective algorithm (Branch et al. 1999). These regressions describe the relationships between the recorded pixel values and the target camera quantum catch for each sensor. The power law regression has the form:

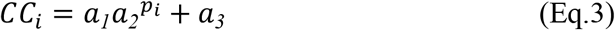

while the polynomial regression has the form.

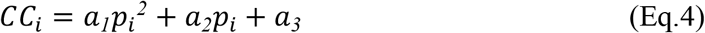

where *p_i_* is the pixel value of the channel *i* and *a*_1_, *a*_2_, *a*_3_ are parameters fit to the data. We then use these linearization parameters to transform all observed pixel values into relative linearized camera responses (relativized such that an object with 100% reflectance would have a value of 1.0). The linearization parameters need to be estimated for each recording session separately and the illumination and camera settings should be maintained while recording images or videos.

Once this step is completed, the camera quantum catches from all four color channels are effectively linearized and equalized. This step removes the effects of unequal illumination from calibration images or frames, provided that the illumination is intense enough across the entire spectrum (Stevens et al. 2007); however, all other frames will be with reference to these calibration frames, even if there are subsequent shifts in illumination. Therefore, these steps should be repeated whenever the illumination changes. Just like other applications of multispectral imagery (Stevens et al. 2007; Troscianko and Stevens 2015), we recommended placing standards in each frame, or recording standards immediately before and/or after a photo session.

### Transformation

Natural spectra take on a limited variety of spectral shapes (Maloney 1986). This regularity enables the mapping of camera responses (pixel values) to photoreceptor quantum catches, which would otherwise be impossible. Thus, we can empirically derive a transformation matrix that reliably transforms from the camera quantum catch to animal quantum catch (Finlayson and Süsstrunk 2000; Westland and Ripamonti 2004; Stevens et al. 2007; Pike 2011). This transformation matrix is specific to the camera sensors, the target animal’s photoreceptor sensitivities, and to the illumination under which the animal photoreceptor responses are to be estimated. It does not however depend on the recording session (Stevens et al. 2007).

When estimating a transformation matrix, first we calculate the camera quantum catches *(CC_i_*, see above) for each object from a large database of spectral reflectance data under ideal illumination. We then calculate the expected photoreceptor quantum catch of the target animal, under any illumination, using the equation:

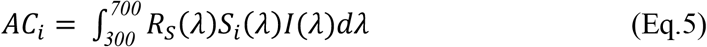

where *AC_i_* is the quantum catch of the photoreceptor *i*, *R_s_* is the spectral reflectance function of the stimulus, *S_i_* is the spectral sensitivity function of the receptor *i*, and *I* is the photon flux of the illuminant. From these two estimates we derive a transformation matrix, *T*, that maps the camera sensor space into the animal receptor space:

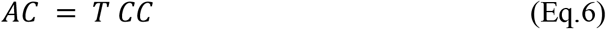

where *AC* is the vector of animal receptor responses and *CC* is the vector of camera quantum catches. The entries of *T* were estimated by fitting linear models (without intercept) such that:

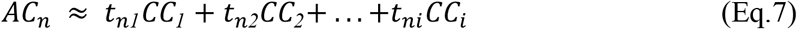

In this paper we used a large database of natural flower reflectance data (FReD, consisting of 2,494 spectra; Arnold et al. 2010) to fit the transformation matrix *T*, for two example animals (the honeybee *Apis mellifera* and the average ultraviolet-sensitive avian viewer (Ender and Mielke 2005), photoreceptor sensitivities downloaded from pavo (Maia et al. 2013), under isoluminant viewing illumination. The predicted animal quantum catches (*AC_n_*) matched the calculated animal quantum catches, for all channels (Figure S13).

### Evaluating the success of the system

To ground-truth our method we compared animal photoreceptor quantum catches calculated directly from reflectance spectra to those derived using our camera system and transformation functions. The photoreceptor quantum catches estimated using the two methods match well (see below).

First, we tested our method on a selection of color standards: an ARUCO standard made from 20 pastels (Figure S8, Table S8) and a custom grayscale made from barium sulfate paint and black 3.0 (see above for details), another set of pastels (Figure S14, Table S8) and a DKK Color Calibration Chart). We repeated the validation tests for stills and videos, under natural, direct and indirect sunlight and in the lab under metal halide illumination (315W, iGrowtek MH315/10K), for the two target animals: honeybees *(Apis mellifera)* and the average ultraviolet-sensitive avian viewer (Ender and Mielke 2005, Maia et al. 2013). In all cases, we found good agreement between the reflectance-based and camera-based estimations (Figures S1-S6, Tables S1-S6, 0.981 < R^2^ < 0.992 for videos in direct sunlight; 0.928 < R^2^ < 0.965 for videos in indirect sunlight; 0.962 < R^2^ < 0.983 for images taken in direct sunlight; 0.928 < R^2^ < 0.961 for images taken in the lab). In addition, we verified that the use of Spectralon standards and a custom-made grayscale provided similar results (Figures S1-S4, Tables S1-S4, linearized on Spectralon 0.942 < R^2^ < 0.985, linearized on ARUCO 0.928 < R^2^ < 0.983, depending on the channel and the lighting).

Next, we looked into the applicability of the method under field conditions. We conducted validation tests on photos of a diverse set of natural objects, including flowers, leaves, birds’ eggs, and feathers (Table S9). Again, for both target animals, photography and spectroscopy provided similar estimates of quantum catches (Figure S7, Table S7, 0.826 < R^2^ < 0.940). As expected, the linear relationship between the two estimates was weaker for natural objects than for color standards. We argue, however, that this mismatch represents a meaningful variation: the perceived color of these objects is altered by their shape (resulting in differences in lighting), fine patterning, texture, and physical color (for iridescent feathers).

We want to stress that the R^2^ reported here and in comparable studies (e.g., Troscianko and Stevens 2015) must be interpreted with caution. The reported coefficient of determination, R^2^, describes the squared correlation coefficient, and only quantifies the strength of the linear relationship between the expected and predicted quantum catches, but it is insensitive to cases where the linear relationship deviates from the 1:1 slope. Thus, this R^2^ does not allow interpretation of the accuracy of the predicted animal quantum catch. Therefore, we report the more informative mean absolute prediction error (MAPE), root mean squared prediction error (RMSPE), as well as the range of the inner 75% of the ordered prediction errors. These three measures are on the same scale as the expected quantum catches and allow for direct assessment of how well the system can predict animal quantum catch.

### Software

The software system was designed with end-users in mind, and it uses automation and interactive platforms where possible. The functionality is packaged as a Python library called video2vision and can be downloaded and installed from the PyPI repository or from github (see Data availability). Operations are grouped into directed acyclic graphs which we call pipelines, essentially a flowchart of operations with no loops (implemented with NetworkX library, Hagberg et al. 2008). Pipelines are saved as JSON files for reuse. Still images or video frames are loaded as numpy arrays (Harris et al. 2020) and processed in batches to minimize runtime. Most image processing operations are performed using the OpenCV library (Bradski 2000). The system can be called from the command line to process one or more images or videos, but a few components that require greater interactivity - such as specifying the spatial relationship between ARUCO markers and sample points - are implemented in Jupyter notebooks (Kluyver et al. 2016). The software system is designed to be easily extendable; new image processing operations can be quickly added as new classes following a defined API.

## Data availability

All data and parts are freely available at figshare (Link available upon acceptance, until then, available upon request), codes are freely available at GitHub and PyPI. Links are provided below.

Github link: https://github.com/hanleycolorlab/video2vision

PyPI link: https://pypi.org/project/video2vision

## Notes

https://github.com/hanleycolorlab/video2vision

https://pypi.org/project/video2vision

https://www.youtube.com/watch?v=dtCx6xDEHRQ

### Competing Interest Statement

The authors have declared no competing interest.

