## Supplementary information for "Recording animal-view videos of the natural world"

**Table S1. Error estimations for prediction for images of known standards, taken under full sunlight and linearized to Spectralon standards**. Here we describe how well our predicted animal quantum catches fit to expected animal quantum catches for the honeybee (*Apis* sp.) and the average UVS avian receiver (Avian). To more fully assess accuracy, we present mean absolute prediction error (MAPE), root mean squared prediction error (RMSPE) as well as the linear association between predicted and expected quantum catch ($R^2$), and the range of the inner 75% of errors (i.e., excluding the 25% largest absolute errors; 75% error band).

| Visual system | Channel | RMSPE | MAPE | $R^2$ | 75% error band |
|---|---|---|---|---|---|
| *Apis* | UV | 0.041 | 0.031 | 0.977 | [-0.044; 0.036] |
| | blue | 0.051 | 0.037 | 0.971 | [-0.058; 0.063] |
| | green | 0.033 | 0.027 | 0.985 | [-0.035; 0.032] |
| Avian | UV | 0.035 | 0.027 | 0.98 | [-0.035; 0.032] |
| | blue | 0.048 | 0.036 | 0.974 | [-0.058; 0.058] |
| | green | 0.066 | 0.047 | 0.965 | [-0.062; 0.051] |
| | red | 0.069 | 0.047 | 0.975 | [-0.059; 0.027] |

**Table S2. Error estimations for prediction for images of known standards, taken under full sunlight and linearized to ARUCO standards.** Here we describe how well our estimated animal quantum catches fit to expected animal quantum catches for the honeybee (*Apis* sp.) and the average UVS avian receiver (Avian). To more fully assess accuracy, we present mean absolute prediction error (MAPE), root mean squared prediction error (RMSPE) as well as the linear association between predicted and expected quantum catch ($R^2$), and the range of the inner 75% of errors (i.e., excluding the 25% largest absolute errors; 75% error band).

| Visual system | Channel | RMSPE | MAPE | $R^2$ | 75% error band |
|---|---|---|---|---|---|
| *Apis* | UV | 0.036 | 0.031 | 0.976 | [-0.039; 0.041] |
| | blue | 0.049 | 0.039 | 0.962 | [-0.055; 0.048] |
| | green | 0.037 | 0.03 | 0.983 | [-0.039; 0.042] |
| Avian | UV | 0.035 | 0.029 | 0.979 | [-0.039; 0.039] |
| | blue | 0.05 | 0.041 | 0.969 | [-0.058; 0.059] |
| | green | 0.07 | 0.05 | 0.967 | [-0.065; 0.070] |
| | red | 0.066 | 0.046 | 0.974 | [-0.054; 0.043] |

**Table S3. Error estimations for prediction for images of known standards, taken under lab light and linearized to Spectralon standards**. Here we describe how well our estimated animal quantum catches fit to expected animal quantum catches for the honeybee (*Apis* sp.) and the average UVS avian receiver (Avian). To more fully assess accuracy, we present mean absolute prediction error (MAPE), root mean squared prediction error (RMSPE) as well as the linear association between predicted and expected quantum catch ($R^2$), and the range of the inner 75% of errors (i.e., excluding the 25% largest absolute errors; 75% error band).

| Visual system | Channel | RMSPE | MAPE | $R^2$ | 75% error band |
|---|---|---|---|---|---|
| *Apis* | UV | 0.058 | 0.043 | 0.944 | [-0.059; 0.058] |
| | blue | 0.07 | 0.05 | 0.942 | [-0.064; 0.068] |
| | green | 0.054 | 0.038 | 0.966 | [-0.049; 0.047] |
| Avian | UV | 0.055 | 0.039 | 0.951 | [-0.044; 0.064] |
| | blue | 0.066 | 0.047 | 0.95 | [-0.055; 0.072] |
| | green | 0.09 | 0.061 | 0.953 | [-0.084; 0.085] |
| | red | 0.073 | 0.051 | 0.953 | [-0.063; 0.067] |

**Table S4. Error estimations for prediction for images of known standards, taken under lab light and linearized to ARUCO standards**. Here we describe how well our estimated animal quantum catches fit to expected animal quantum catches for the honeybee (*Apis* sp.) and the average UVS avian receiver (Avian). To more fully assess accuracy, we present mean absolute prediction error (MAPE), root mean squared prediction error (RMSPE) as well as the linear association between predicted and expected quantum catch ($R^2$), and the range of the inner 75% of errors (i.e., excluding the 25% largest absolute errors; 75% error band).

| Visual system | Channel | RMSPE | MAPE | $R^2$ | 75% error band |
|---|---|---|---|---|---|
| *Apis* | UV | 0.055 | 0.041 | 0.929 | [-0.062; 0.056] |
| | blue | 0.071 | 0.053 | 0.928 | [-0.069; 0.071] |
| | green | 0.059 | 0.042 | 0.961 | [-0.049; 0.059] |
| Avian | UV | 0.059 | 0.042 | 0.934 | [-0.065; 0.043] |
| | blue | 0.066 | 0.05 | 0.943 | [-0.065; 0.066] |
| | green | 0.102 | 0.071 | 0.948 | [-0.075; 0.088] |
| | red | 0.082 | 0.057 | 0.94 | [-0.064; 0.074] |

**Table S5. Error estimations for prediction for videos of known standards, taken under full sunlight and linearized to Spectralon standards**. Here we describe how well our estimated animal quantum catches fit to expected animal quantum catches for the honeybee (*Apis* sp.) and the average UVS avian receiver (Avian). To more fully assess accuracy, we present mean absolute prediction error (MAPE), root mean squared prediction error (RMSPE) as well as the linear association between predicted and expected quantum catch ($R^2$), and the range of the inner 75% of errors (i.e., excluding the 25% largest absolute errors; 75% error band).

| Visual system | Channel | RMSPE | MAPE | $R^2$ | 75% error band |
|---|---|---|---|---|---|
| *Apis* | UV | 0.036 | 0.028 | 0.982 | [-0.037; 0.044] |
| | blue | 0.041 | 0.028 | 0.985 | [-0.031; 0.027] |
| | green | 0.038 | 0.026 | 0.99 | [-0.022; 0.029] |
| Avian | UV | 0.038 | 0.028 | 0.981 | [-0.044; 0.010] |
| | blue | 0.042 | 0.027 | 0.989 | [-0.024; 0.036] |
| | green | 0.043 | 0.029 | 0.992 | [-0.017; 0.038] |
| | red | 0.06 | 0.041 | 0.988 | [-0.022; 0.065] |

**Table S6. Error estimations for prediction for videos of known standards, taken in shade, under sunlight and linearized to ARUCO standards**. Here we describe how well our estimated animal quantum catches fit to expected animal quantum catches for the honeybee (*Apis* sp.) and the average UVS avian receiver (Avian). To more fully assess accuracy, we present mean absolute prediction error (MAPE), root mean squared prediction error (RMSPE) as well as the linear association between predicted and expected quantum catch ($R^2$), and the range of the inner 75% of errors (i.e., excluding the 25% largest absolute errors; 75% error band).

| Visual system | Channel | RMSPE | MAPE | $R^2$ | 75% error band |
|---|---|---|---|---|---|
| *Apis* | UV | 0.074 | 0.058 | 0.934 | [-0.042; 0.098] |
| | blue | 0.087 | 0.067 | 0.928 | [-0.045; 0.096] |
| | green | 0.079 | 0.063 | 0.943 | [-0.076; 0.091] |
| Avian | UV | 0.085 | 0.068 | 0.935 | [-0.025; 0.101] |
| | blue | 0.081 | 0.061 | 0.936 | [-0.077; 0.093] |
| | green | 0.111 | 0.089 | 0.933 | [-0.108; 0.121] |
| | red | 0.069 | 0.055 | 0.965 | [-0.073; 0.076] |

**Table S7. Error estimations for prediction for images of natural objects, taken under full sunlight and linearized to ARUCO standards.** Here we describe how well our estimated animal quantum catches fit to expected animal quantum catches for the honeybee (*Apis* sp.) and the average UVS avian receiver (Avian). To more fully assess accuracy, we present mean absolute prediction error (MAPE), root mean squared prediction error (RMSPE) as well as the linear association between predicted and expected quantum catch ($R^2$), and the range of the inner 75% of errors (i.e., excluding the 25% largest absolute errors; 75% error band).

| **Visual system** | **Channel** | **RMSPE** | **MAPE** | **$R^2$** | **75% error band** |
|---|---|---|---|---|---|
| *Apis* | UV | 0.043 | 0.028 | 0.938 | [-0.025; 0.032] |
| | blue | 0.05 | 0.036 | 0.917 | [-0.055; 0.054] |
| | green | 0.07 | 0.05 | 0.877 | [-0.069; 0.067] |
| Avian | UV | 0.041 | 0.027 | 0.94 | [-0.029; 0.031] |
| | blue | 0.058 | 0.042 | 0.91 | [-0.055; 0.056] |
| | green | 0.101 | 0.069 | 0.851 | [-0.077; 0.091] |
| | red | 0.096 | 0.07 | 0.826 | [-0.097; 0.089] |

**Table S8 Custom color cards.** Here we provide information on the two custom color cards that we used in this study. The first was the custom color card with ARUCO markers, which includes 20 pastels and 8 grayscale patches (see Methods and Materials for details). The second was a small color card, visible in a few shots (e.g., Figure S14). We prefix each color target with a number corresponding with its position on the card (see Figure S8, S14) and its position within the reflectance spectra dataset.

| Standard | Material | Manufacturer descriptions | Target description |
|---|---|---|---|
| ARUCO standard | Blick Artists' Soft Pastel | 20 Artists' Pastels Half Sticks. 21948-1209 | (1) Black[1], (2) White[1], (3) Cool Grey 3[1], (4) Burnt Umber 4, (5) Burnt Sienna 4, (6) Raw Sienna 2 |
| | Barium sulfate paint and black paint | Labsphere 6080, and Culture Hustle, Black 3.0 | (7) ~99%, (8) ~70%, (9) ~50%, (10) ~43%, (11) ~16%, (12) ~11%, (13) ~5%, (14) ~3% |
| | Blick Artists' Soft Pastel | 20 Artists' Pastels Half Sticks. 21948-1209 | (15) Yellow Ochre 4, (16) Olive Green 4[2], (17) Sap Green 2, (18) Viridian Hue 4[2], (19) Phthalo Blue 3 (Green Shade), (20) Prussian Blue 1, (21) Ultramarine 4, (22) Purple 3, (23) Crimson Lake 1, (24) Cadmium Red Hue 4, (25) Cadmium Red Orange Hue 4, (26) Cadmium Orange Hue 2, (27) Yellow 3, (28) Lemon Yellow 2 |
| Pastel card | Blick Artists' Soft Pastel | 20079-XXXX[3] | (1) White[1], (2) Cool gray 1, (3) Sepia 1, (4) Cool gray 2, (5) Cool gray 3[1], (6) Cool gray 4, (7) Blue gray 4, (8) Black[1], (9) Phthalo blue (red shade) 4, (10) Lemon yellow 4, (11) Bright green 4, (12) Crimson lake 4, (13) Purple brown 1 |

[1] This color was present in both color cards, though validation tests were conducted using reflectance spectra from each sample.

[2] This color did not properly adhere to the surface and was excluded from tests.

[3] These colors were purchased individually, and each would have a different serial number in the 20079 series.

**Table S9 Natural objects tested.** Here we provide information on the natural objects we used in this study, indicating the region and how that region was coded on spectral reflectance measurements.

| **Common name** | **Species** | **Specimen type** | **Regions/ Measurement IDs** |
|---|---|---|---|
| Showy goldeneye | *Heliomeris multiflora* | flower | tip, base |
| Marsh mallow | *Althaea officinalis* | flower | tip, base |
| Black-eyed Susan | *Rudbeckia hirta* | flower | tip base |
| Maximilian sunflower | *Helianthus maximiliani* | flower | tip, base |
| Fameflower | *Talinum paniculatum* | flower | tip, base |
| Everlasting pea | *Lathyrus latifoliés* | flower | tip |
| Burning bush | *Euonymus alatus* | leaf | leaf 1, leaf 4, leaf 8, leaf 14, leaf 19, leaf 23, leaf 25, leaf 29, leaf 32, leaf 36 |
| Shadblow serviceberry | *Amelanchier canadensis* | leaf | leaf 2, leaf 10, leaf 13, leaf 31, leaf 34 |
| Eastern black walnut | *Juglans nigra* | leaf | leaf 3, leaf 12 |
| American bittersweet | *Celastrus scandens* | leaf | leaf 5 |
| Golden raintree | *Koelreuteria paniculata* | leaf | leaf 6, leaf 7, leaf 18, leaf 26, leaf 37 |
| Pignut Hickory | *Carya glabra* | leaf | leaf 9, leaf 16, leaf 21, leaf 22, leaf 33, leaf 35 |
| White oak | *Quercus alba* | leaf | leaf 11 (back) |
| Northern red oak | *Quercus rubra* | leaf | leaf 17, leaf 20 |
| Victoria creeper | *Parthenocissus quinquefolia* | leaf | leaf 27 |
| Fox grape | *Vitis vulpina* | leaf | leaf 28, leaf 39 |
| Hackberry | *Prunus padus* | leaf | leaf 30 |
| Bur oak | *Quercus macrocarpa* | leaf | leaf 38 |
| Black locust | *Robinia pseudoacacia* | leaf | leaf 40 |
| Japanese quail | *Coturnix coturnix* | egg[1] | egg A ground, egg A spot, egg A spot, egg B ground |
| American robin | *Turdus migratorius* | egg[2] | egg A ground, egg B ground |
| Gray catbird | *Dumetella carolinensis* | egg[2] | egg A ground, egg B ground |
| Brown thrasher | *Toxostoma rufum* | egg[2] | egg A ground, egg B ground, egg C ground |
| Northern mockingbird | *Mimus polyglottos* | egg[2] | egg A ground with spots mixed, egg A blue ground, egg A spot, egg B ground, egg C ground |

| **Common name** | **Species** | **Specimen type** | **Regions/ Measurement IDs** |
|---|---|---|---|
| Mourning dove | *Zenaida macroura* | egg[2] | egg A ground, egg B ground |
| Domestic duck | *Anas platyrhynchos* | egg[1] | egg A, egg B |
| Baltimore oriole | *Icterus galbula* | bird[3] | breast, belly |
| American Kestrel | *Falco sparverius* | bird[3] | breast, throat, spot |
| Eastern bluebird | *Sialia sialis* | bird[3] | breast, belly, rump |
| Northern cardinal | *Cardinalis cardinalis*, male | bird[3] | breast, throat |
| Blackburnian warbler | *Setophaga fusca,* male | bird[3] | throat, belly, black streak |
| Brown-headed cowbird | *Molothrus ater*, female | bird[3] | breast |
| Indigo bunting | *Passerina cyanea,* male | bird[3] | belly, breast, throat |
| American goldfinch | *Spinus tristis,* male | bird[3] | breast |
| Banaquit | *Coereba flaveola* | bird[3] | belly, breast, throat |
| Ruby-throated hummingbird | *Archilocus colubris,* male | bird[3] | tail, breast, throat |
| European starling | *Sturnus vulgaris* | bird[3] | belly (green area), throat |

1. Commercially purchased.
2. Abandoned egg collected under US Fish and Wildlife Service Collecting permit (MB81216C-3) and Virginia Department of Game and Inland Fisheries Scientific collecting permit (070605)
3. Provided by the teaching collection at George Mason University

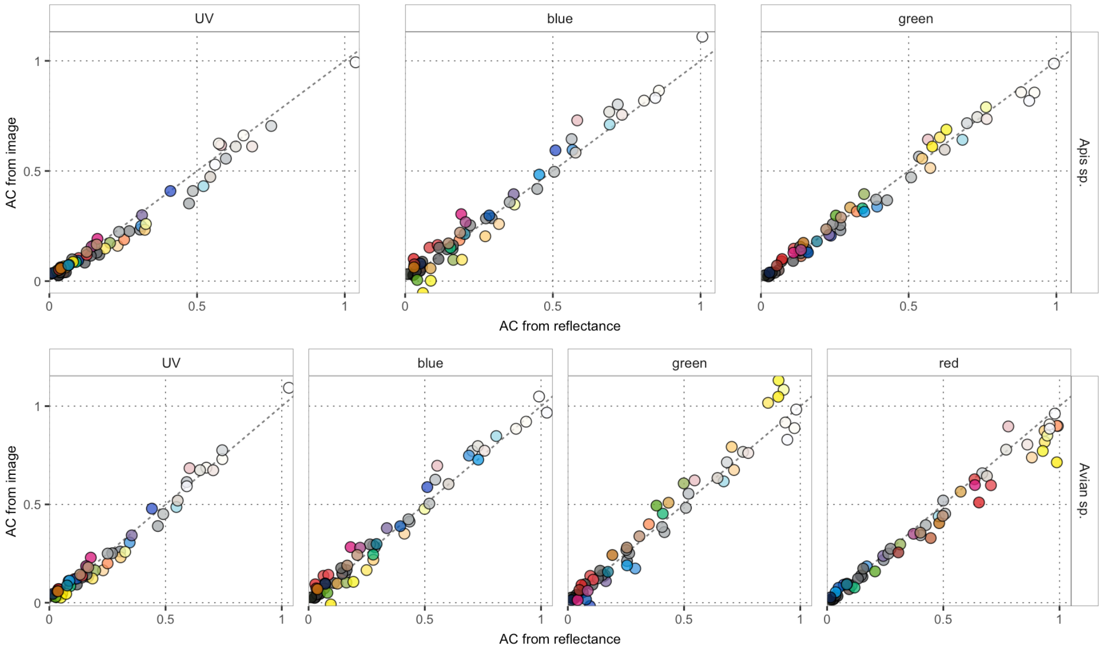


**Figure S1. Evaluating the fit for images of known standards.** In this case, the images were taken under full sunlight and linearized to a set of Spectralon standards. The plots show the expected animal quantum catch (AC from reflectance) against our predicted animal quantum catch (AC from image). We plot the fit for both the honeybee (*Apis* sp., top) and the average ultraviolet sensitive avian receiver (*Avian* sp., bottom), for each of their three and four photoreceptors, respectively. The marker colors indicate the human-perceived color of the sample. For data on fit, please see Table S1.

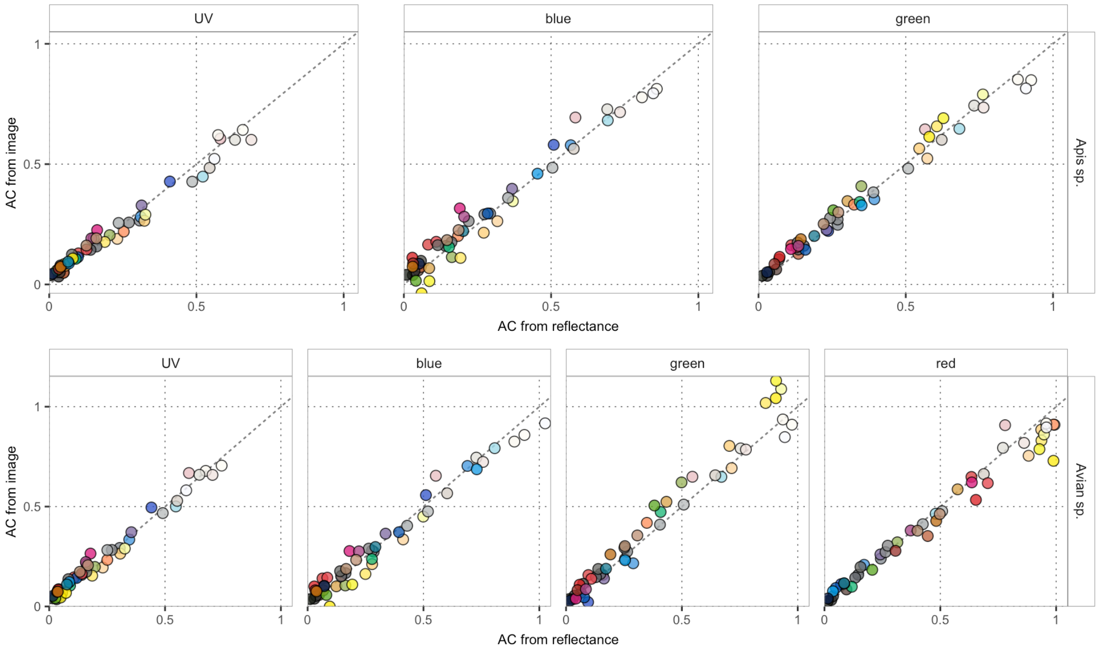


**Figure S2. Evaluating the fit for images of known standards.** In this case, the images were taken under full sunlight and linearized to a set of ARUCO standards. The plots show the expected animal quantum catch (AC from reflectance) against our predicted animal quantum catch (AC from image). We plot the fit for known color standards, and for both the honeybee (*Apis* sp., top) and the average ultraviolet sensitive avian receiver (*Avian* sp., bottom), for each of their three and four photoreceptors, respectively. The marker colors indicate the human-perceived color of the sample. For data on fit, please see Table S2.

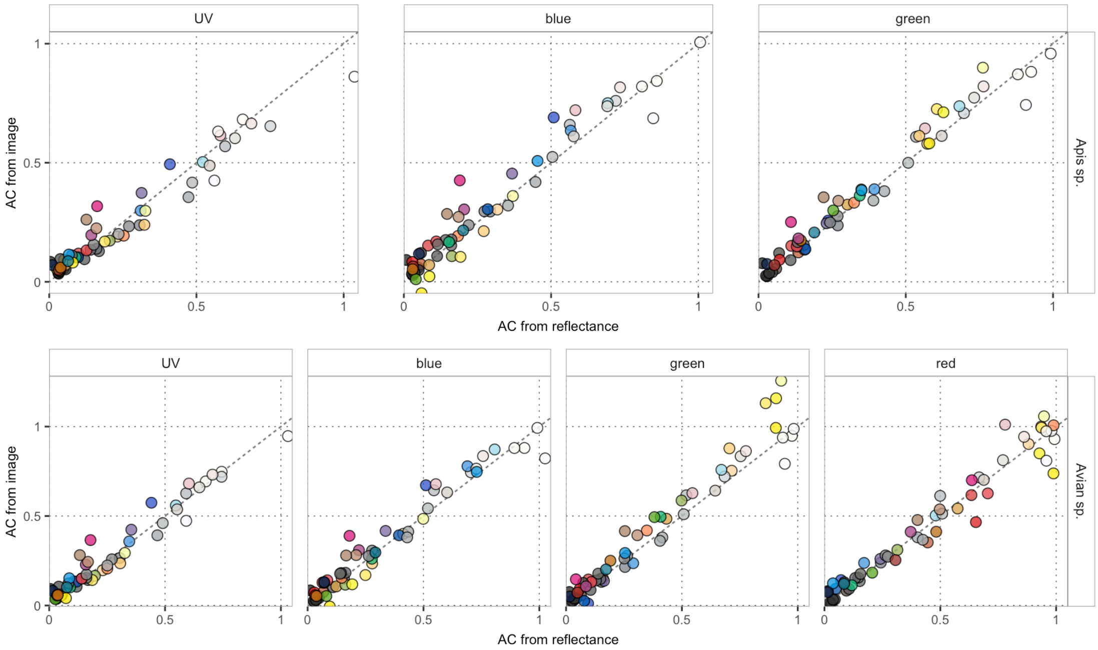


**Figure S3. Evaluating the fit for images of known standards.** In this case, the images were taken under lab light and linearized to a set of Spectralon standards. The plots show the expected animal quantum catch (AC from reflectance) against our predicted animal quantum catch (AC from image). We plot the fit for known color standards, and for both the honeybee (*Apis* sp., top) and the average ultraviolet sensitive avian receiver (*Avian* sp., bottom), for each of their three and four photoreceptors, respectively. The marker colors indicate the human-perceived color of the sample. For data on fit, please see Table S3.

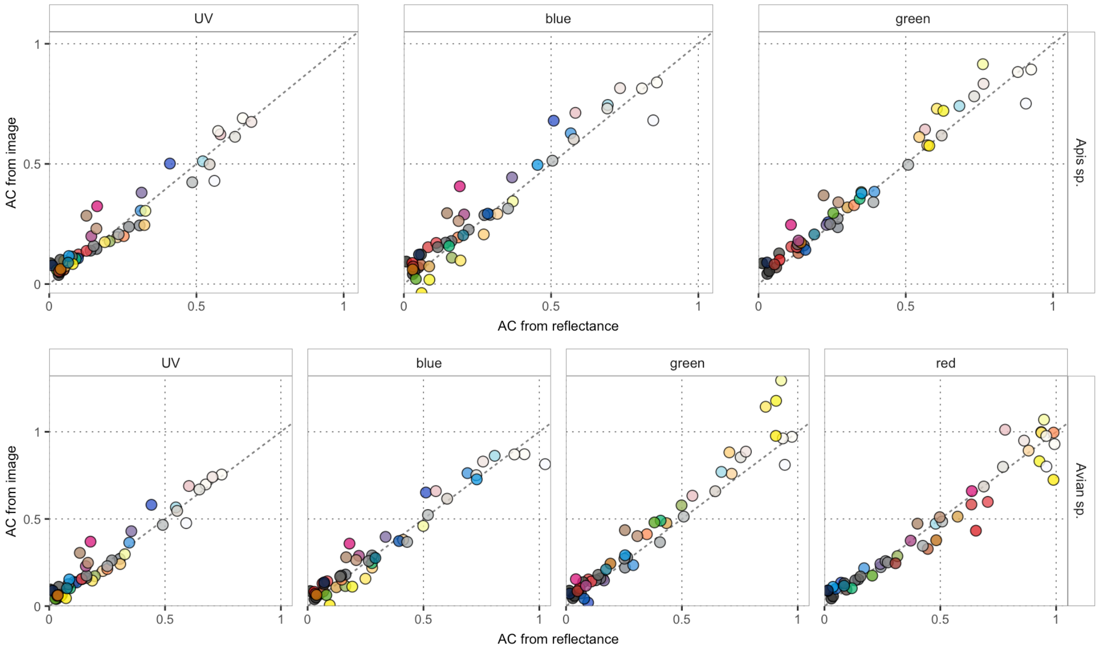


**Figure S4. Evaluating the fit for images of known standards.** In this case, the images were taken under lab light and linearized to a set of ARUCO standards. The plots show the expected animal quantum catch (AC from reflectance) against our predicted animal quantum catch (AC from image). We plot the fit for known color standards, and for both the honeybee (*Apis* sp., top) and the average ultraviolet sensitive avian receiver (*Avian* sp., bottom), for each of their three and four photoreceptors, respectively. The marker colors indicate the human-perceived color of the sample. For data on fit, please see Table S4.

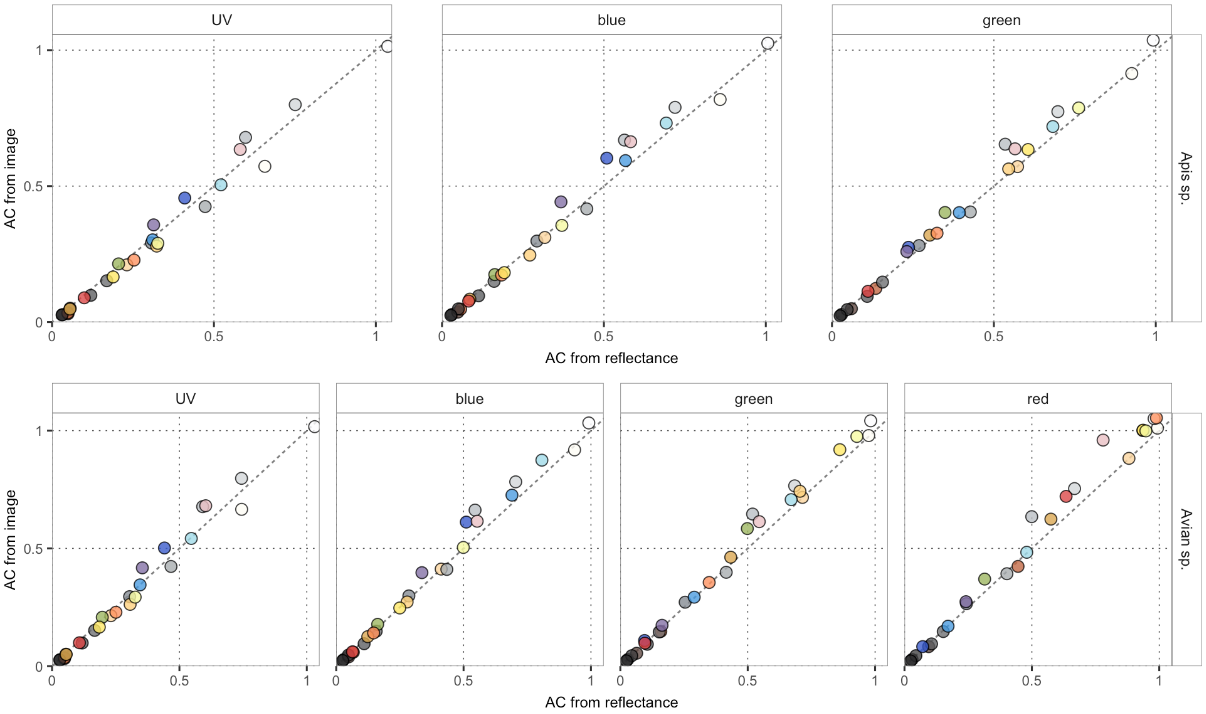


**Figure S5. Evaluating the fit for videos of known standards.** In this case, the videos were taken under full sunlight and linearized to a set of Spectralon standards. The plots show the expected animal quantum catch (AC from reflectance) against our predicted animal quantum catch (AC from image). We plot the fit for known color standards, and for both the honeybee (*Apis* sp., top) and the average ultraviolet sensitive avian receiver (*Avian* sp., bottom), for each of their three and four photoreceptors, respectively. The marker colors indicate the human-perceived color of the sample. For data on fit, please see Table S5.

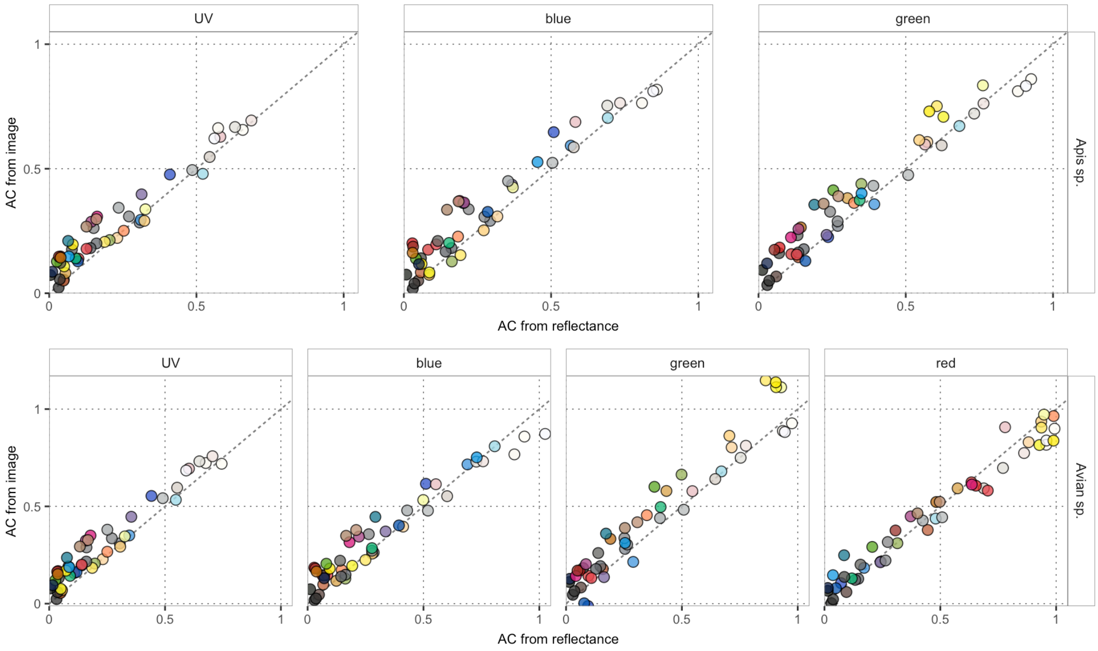


**Figure S6. Evaluating the fit for videos of known standards.** In this case, the videos were taken in shade, under sunlight, and linearized to a set of ARUCO standards. The plots show the expected animal quantum catch (AC from reflectance) against our predicted animal quantum catch (AC from image). We plot the fit for known color standards, and for both the honeybee (*Apis* sp., top) and the average ultraviolet sensitive avian receiver (*Avian* sp., bottom), for each of their three and four photoreceptors, respectively. The marker colors indicate the human-perceived color of the sample. For data on fit, please see Table S6.

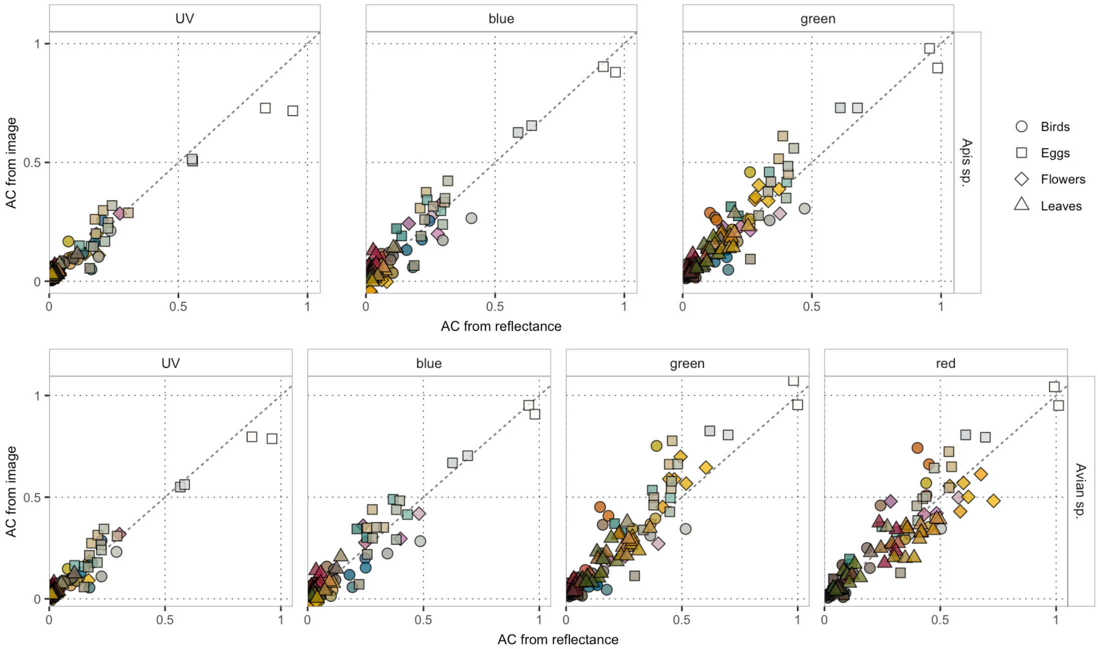


**Figure S7. Evaluating the fit of natural objects**. In this case, the images were taken under full sunlight and linearized to a set of ARUCO standards. The plots show the expected animal quantum catch (AC from reflectance) against our predicted animal quantum catch (AC from images). We plot the fit for a collection of flowers (diamonds), leaves (triangles), birds' eggs (squares) and birds' feathers (circles); see Table S9 for sample details. The fit is shown for both the honeybee (*Apis* sp., top) and the average ultraviolet sensitive avian receiver (*Avian* sp., bottom), for each of their three and four photoreceptors, respectively. The linear relationship between the two estimates was weaker for natural objects than for color standards. The mismatch represents a meaningful variation: the perceived color of natural objects is altered by their shape, fine patterning, texture, and physical color. The marker colors indicate the human-perceived color of the sample. For data on fit, please see Table S7.

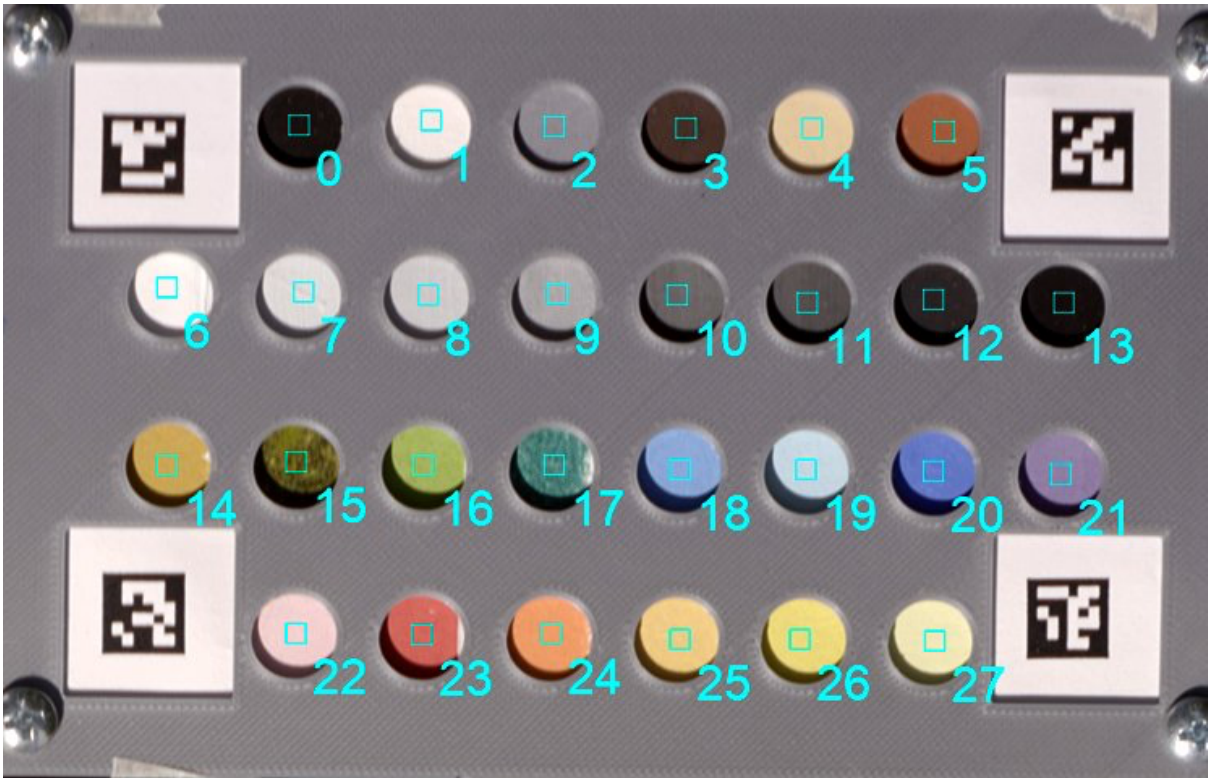


**Figure S8. A custom color standard.** We used a custom color card that featured four ARUCO fiducial markers and 28 distinct colors. The color patches were made from pastels (positions 0-5 & 14-27) or a mixture of barium sulfate (white) and flat black (Black 3.0) paints (positions 6-13). These provide fast, highly accurate, automated methods for calibration, linearization, and transformation. For details on how to construct these cards, see Table S8.

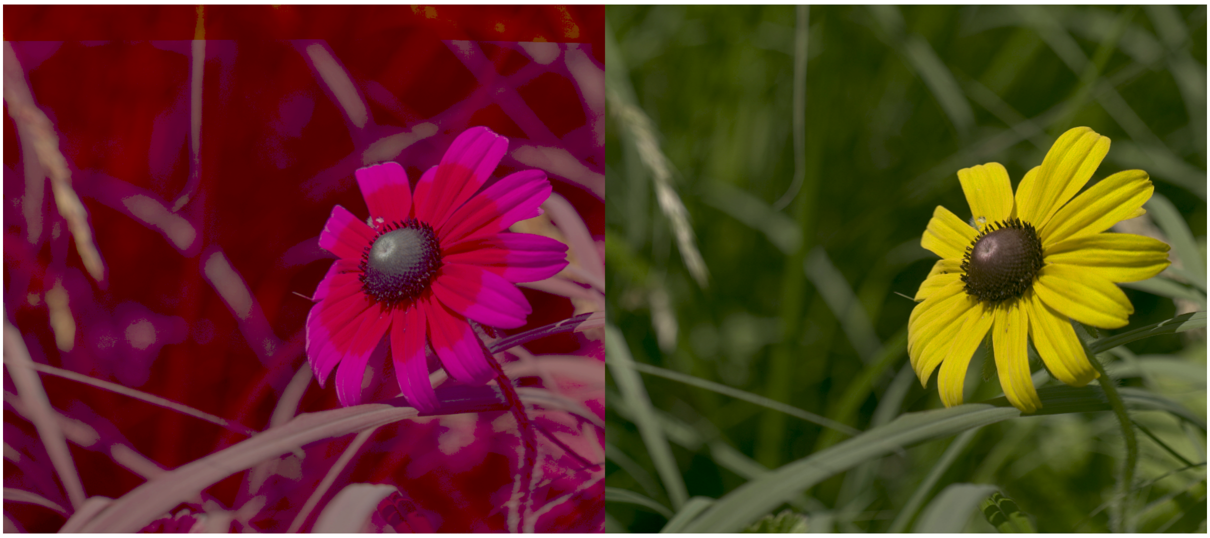

**Figure S9.** A black-eyed Susan (*Rudbeckia hirta*) depicted as a honeybee (*Apis mellifera*) false color image and the same flower as a linear, human-vision, image. We have used a gamma correction on the honeybee false color and linear image ($AC_i^{0.3}$ and $CC_i^{0.5}$, respectively) to enhance contrast. These are the same images as shown in Figure 1.

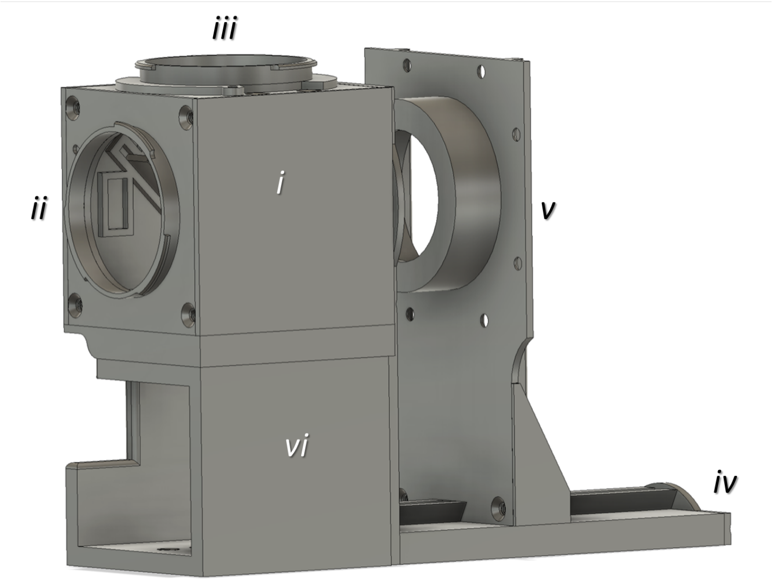


**Figure S10. Modular 3D printed housing.** A schematic showing the (i) cage, with mounting points for the (ii) visible light Sony camera, the (iii) full spectrum modified camera, a (iv) rail to allow for focusing, a (v) lens and bag bellows attachment plate, and a (vi) spacer to facilitate various shooting angles while tripod mounted.

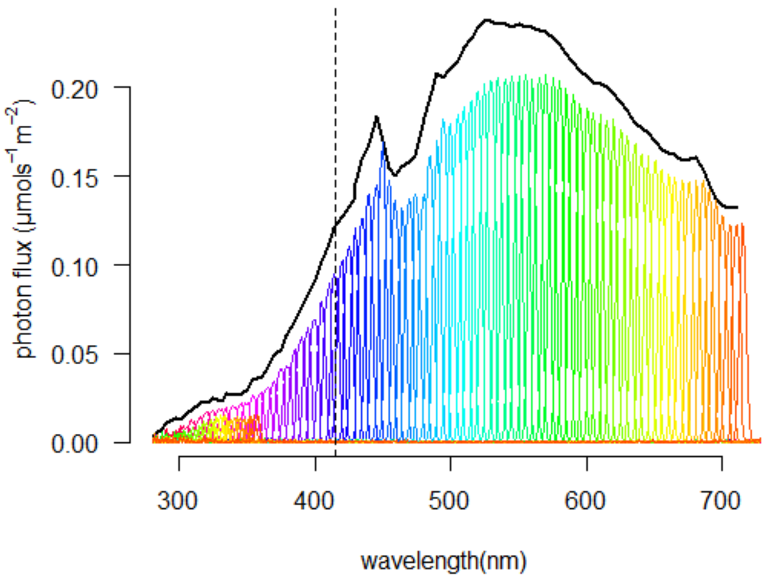


**Figure S11. Illumination used for estimating sensor sensitivities.** We used a monochromator to deliver narrow bands of light (colored bands, mean FWHM ± s.e. = 7.3 ± 0.29 nm) from 280-800 nm cosine corrector that was made of white Spectralon, which was then imaged. The total power used for estimation (solid black line) was the sum photo flux (μmol $s^{-1}$ $m^{-2}$).

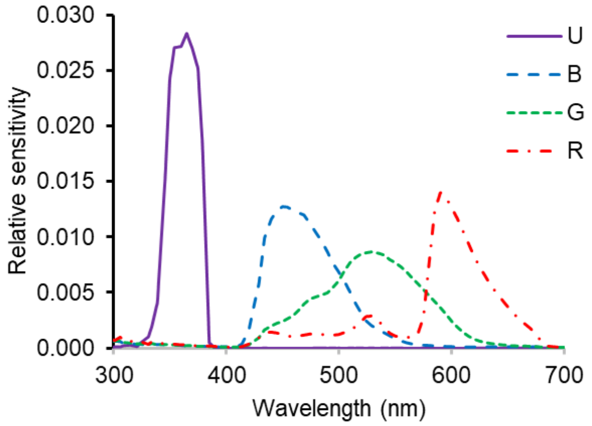


**Figure S12. Camera sensor sensitivity.** Estimates of relative camera sensor sensitivity for the ultraviolet (U, solid purple line), blue (B, dashed blue line), green (G, dotted green line), and red (R, dot-dashed red line) sensors.

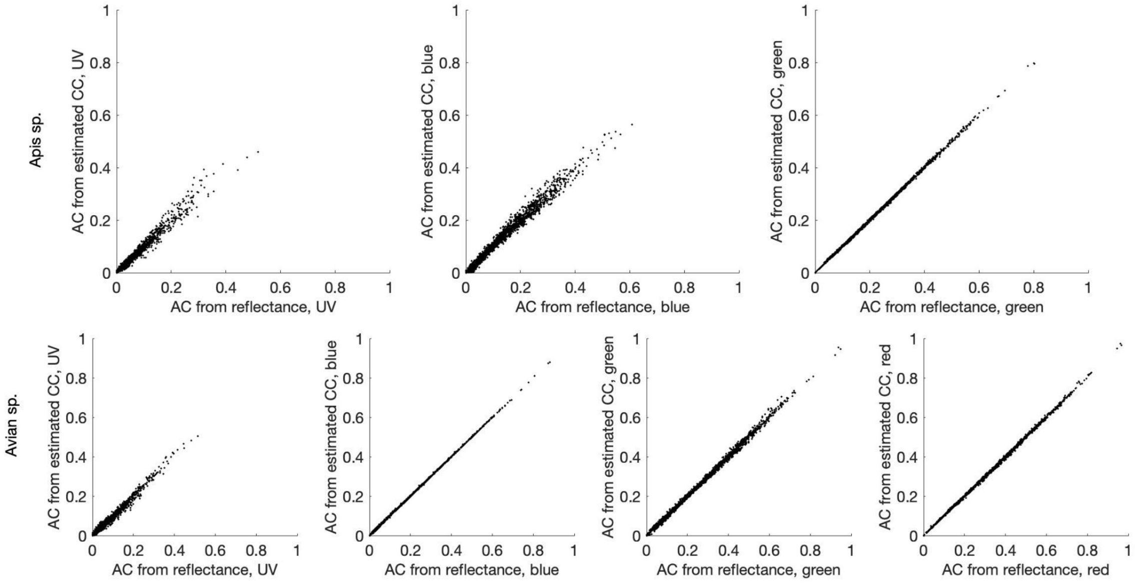


**Figure S13. Relationship between the expected and predicted animal quantum catches for the training library.** In this paper we used 2,494 spectra from the FReD database (Arnold et al. 2010) to derive a transformation matrix, *T*, to convert linear camera catches into animal catches. Here we illustrate the fit for those relationships. The models for the blue, green, and red bands for *Avian* sp. and the green band for *Apis* sp. achieve $R^2 \geq 0.998$, while the model fits for the other bands yield an $R^2$ between 0.968 and 0.981. The ranges of the inner 75% of errors across all bands are very narrow: [-0.006; 0.007] for *Apis* sp. and [-0.005; 0.005] for *Avian* sp.

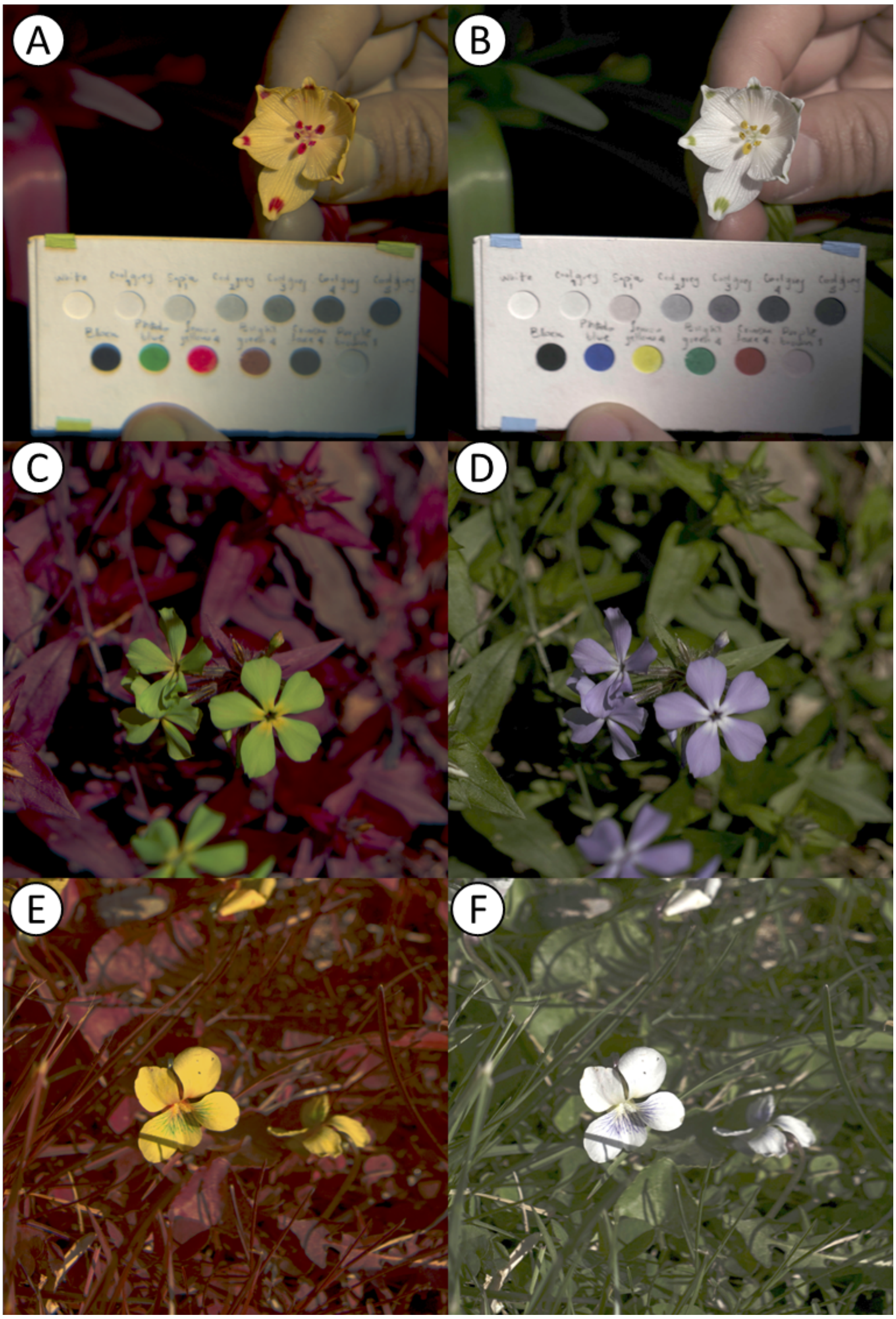


**Figure S14 Examples of honeybee false color images**. Here we illustrate a A-B) Summer snowflake *Leucojum aestivum*, C-D) Blue phlox *Phlox divaricata*, and a D-E) Blue Violet *Viola sororia* in honeybee false color (left) and human visible. We also show a simple, cheap, pastel-based color standard that we used to validate animal-perceived quantum catches (A, B). We applied a gamma correction to this image ($AC_i^{0.3}$ and $CC_i^{0.5}$, respectively).
